# Oxidative Stress Responses and Nutrient Starvation in MCHM Treated *Saccharomyces cerevisiae*

**DOI:** 10.1101/2020.08.17.253799

**Authors:** Michael C. Ayers, Zachary N. Sherman, Jennifer E.G. Gallagher

**Affiliations:** Department of Biology, West Virginia University, Morgantown, WV 26506

**Keywords:** environmental stress response, MCHM, *Saccharomyces cerevisiae*, genetic screen, reactive oxygen species, nutrient starvation, aromatic amino acids, tryptophan

## Abstract

In 2014, the coal cleaning chemical 4-methylcyclohexane methanol (MCHM) spilled into the water supply for 300,000 West Virginians. Initial toxicology tests showed relatively mild results, but the underlying effects on cellular biology were underexplored. Treated wildtype yeast cells grew poorly, but there was only a small decrease in cell viability. Cell cycle analysis revealed an absence of cells in S phase within thirty minutes of treatment. Cells accumulated in G1 over a six-hour time course, indicating arrest instead of death. A genetic screen of the haploid knockout collection revealed 329 high confidence genes required for optimal growth in MCHM. These genes encode three major cell processes: mitochondrial gene expression/translation, the vacuolar ATPase, and aromatic amino acid biosynthesis. The transcriptome showed an upregulation of pleiotropic drug response genes and amino acid biosynthetic genes and downregulation in ribosome biosynthesis. Analysis of these datasets pointed to environmental stress response activation upon treatment. Overlap in datasets included the aromatic amino acid genes *ARO1*, *ARO3*, and four of the five *TRP* genes. This implicated nutrient deprivation as the signal for stress response. Excess supplementation of nutrients and amino acids did not improve growth on MCHM, so the source of nutrient deprivation signal is still unclear. Reactive oxygen species and DNA damage were directly detected with MCHM treatment, but timepoints showed these accumulated slower than cells arrested. We propose that wildtype cells arrest from nutrient deprivation and survive, accumulating oxidative damage through the implementation of robust environmental stress responses.

## Introduction

The chemical 4-methylcyclohexane methanol (MCHM) was a previously little-studied chemical involved in the processing of coal, until a rusted storage tank resulted in a spill of crude MCHM into the Elk River near Charleston, WV. The spill’s size and location adjacent to a drinking water treatment intake were sufficient to fill homes with an overpowering odor that left many fearful of health consequences (Thomasson et al., 2017). This spill interrupted the water supply of approximately 300,000 residents. Research on MCHM since the Elk River spill has increased dramatically over concerns about the lack of characterized physical and biological properties of this chemical (Weidhaas et al., 2016). For instance, an improved toxicological study on MCHM’s effect on model organism viability has been performed (*West Virginia Chemical Spill: Collective NTP Findings and Supporting Files*, n.d.), as well as one study on the potential stress responses it may produce in yeast (Lan et al., 2015). However, these studies had a focus on a few specific toxicological outcomes and predetermined stress pathways that may miss other cellular changes. Recently, research has been begun to identify the biochemical and transcriptional changes that MCHM may produce in organisms (Pupo, Ayers, et al., 2019; Pupo, Ku, et al., 2019).

The environmental stress response (ESR) is a shared transcriptional response to multiple stressors, including heat shock, osmotic shock, and hydrogen peroxide treatment, among others (Gasch et al., 2000). The role of paralogous transcription factors Msn2 and Msn4 are important for a large portion of the ESR transcriptional induction (Gorner et al., 1998; Martínez-Pastor et al., 1996). The transcriptional programming of the environmental stress response incorporates signals from diverse stress pathways including nutrient starvation (TOR), osmolarity stress (HOG), and others, into the Msn2/4 transcriptional activators (Capaldi et al., 2008; De Wever et al., 2005; Garmendia-Torres et al., 2007; Gorner et al., 1998; Gutin et al., 2015; Santhanam et al., 2004). Implementation of the stress response involves two waves of Msn2/4 import to the nucleus that modulate the initial intensity and prolonged duration of the response (Gutin et al., 2019). The signaling kinase Mck1 is indispensable for the prolonged response requiring the second import of active Msn2/4 to the nucleus (Gutin et al., 2019), although it is unclear how the signal is maintained as there does not appear to be direct Mck1 interaction with Msn2 (Hirata et al., 2002). The karyopherin Msn5 involved in nuclear import and export activity is important to maintain the prolonged stress response activity of Msn2/4 in the ESR, apparently through the export of initial wave Msn2 from the nucleus, allowing for the second wave of transcriptional activators to function (Gutin et al., 2019).

Amino acid biosynthesis is predominately regulated by the general amino acid transcriptional activator Gcn4. Under amino acid starvation, the levels of Gcn4 increase due to the decreased degradation and increased translation of the protein (reviewed in Hinnebusch, 1997; Meimoun et al., 2000). Gcn4 binds to a consensus promoter sequence upstream of many of the amino acid biosynthetic genes to positively regulate their expression (Arndt & Fink, 1986; Hill et al., 1986; Oliphant et al., 1989). This amino acid transcriptional control includes the aromatic amino acid biosynthetic gene *ARO3* and the specific tryptophan biosynthetic genes *TRP2, TRP3, TRP4,* and *TRP5* (Braus, 1991). Tryptophan has been implicated for roles in stress tolerances other than nutrient starvation. SDS sensitivity, a cell wall and plasma membrane integrity stress that involves Mck1 effectors in yeast, is dependent on tryptophan biosynthesis and levels of tryptophan and tyrosine in the media and cell (Schroeder & Ikui, 2019). Tryptophan biosynthetic mutants, *trp1-5*, were also reported in a special warning for their use as auxotrophic markers in yeast genetics due to an aberrant sensitivity to many stressors including rapamycin, high pH, and several metal cations (González et al., 2008). A more recent report has shown that tryptophan depletion due to a combination of transporter dysfunction at low temperature and *trp1-5* mutants also confers sensitivity to the DNA damaging agents MMS and HU (Godin et al., 2016). The role of the aromatic amino acids and their biosynthetic pathways in stress tolerance other than nutrient starvation is not fully understood, but it is well established.

Reactive oxygen species (ROS) have previously been implicated as a source of toxicity in cells treated with MCHM (Lan et al., 2015). Cells contain conserved robust networks to mitigate the toxic effects of ROS. These include various proteins, from enzymes that detoxify the reactive species directly, to proteins that repair damage within the cell, such as to DNA (Ayer et al., 2014). The thioredoxin and glutathione (GSH) pathways have significant roles in the cell’s response to ROS. They perform overlapping functions reducing thiol oxidation that can damage proteins in the cytosol. Furthermore, GSH has roles in iron homeostasis between the mitochondria and vacuole, and potentially as a possible buffer for oxidation in disulfide bond formation during protein folding in the ER (Cuozzo & Kaiser, 1999; reviewed by Toledano et al., 2013). Mitochondria serve as a major producer of ROS in the cell as oxidative phosphorylation leaks electrons to molecular oxygen to produce superoxide anions, so these pathways are activated by normal cellular metabolism (reviewed by all the following, Ayer et al., 2014; Perrone et al., 2008; Temple et al., 2005). However, they also become important during the response to toxic chemicals, which can produce ROS directly, or else indirectly through metabolism and attempted detoxification in the vacuole. Toxicity of chemicals that produce ROS in the cell may be mitigated through treatment with antioxidants or intensified through damage to the cellular stress networks (Couto et al., 2016; Sekito et al., 2014).

Much of a yeast cell’s response to stress involves the vacuole (Li & Kane, 2010). This organelle serves as a site for various processes of degradation, detoxification, and metal ion and pH homeostasis (Li & Kane, 2010). The vacuolar ATPase (v-ATPase) is a structure in yeast that is highly conserved and has been adapted to perform a wide range of functions in various eukaryotes. In many animals, including *Drosophila*, homologs of the v-ATPase are known to contribute to nerve function via vesicular excretion of neurotransmitters in animals including humans (Hiesinger et al., 2005). In yeast, the v-ATPase is responsible for acidifying the vacuole interior, creating a proton gradient that is responsible for multiple homeostatic processes (reviewed by the following, N. Nelson & Harvey, 1999; Nishi & Forgac, 2002). As such, its effects on metal ion transport and vacuolar acidification result in a set of knockout phenotypes for many of the v-ATPase’s subunits (Hemenway et al., 1995; H. Nelson & Nelson, 1990; Ohya et al., 1991; Sambade et al., 2005). This *vma-* phenotype includes sensitivities to metal ions, reactive oxygen species, and pH perturbations in either direction. Inositol depletion through mutation or chemical treatment negatively impacts the assembly and activity of the v-ATPase in yeast (Deranieh et al., 2015; Ohya et al., 1991). Any chemical that can inhibit or damage the v-ATPase would likely have distinct consequences for the cell’s ability to cope with other stresses.

The goal of this study was to characterize the response of yeast cells to the foreign chemical MCHM. Treatment of yeast with MCHM created pleiotropic effects on networks throughout the cell. We employed methods including viability assays, RNAseq, flow cytometry, and a genetic screen of knockout strains to characterize these effects. We found several expected changes to networks, such as the pleiotropic drug response ABC transporters that remove xenobiotics from the cell. Our data show direct evidence of ROS and DNA damage following MCHM treatment. The stress also revealed a role for the aromatic amino acid biosynthetic pathway outside nutrient availability for the response to this particular stressor. Any toxicity that these cellular changes caused did not result in large changes in cellular viability but instead resulted in the arrest of the cell cycle in G1. Effects on the cell were wide-ranging, but wildtype cells were able to implement the ESR to recover from exposure at levels higher than spill levels.

## Materials and Methods

### Yeast strains and media

The haploid BY4741 strain (MATA, *his3Δ, leu2Δ, ura3Δ, met15Δ)* (Brachmann et al., 1998) and its MATalpha counterpart BY4742 (*his3Δ, leu2Δ, ura3Δ, lys2Δ*), were used for the majority of experiments as denoted. RNAseq, viability, growth, comet and flow cytometry assays used the BY4741 strain. The genetic screen used the BY4742 collection (Giaever et al., 2002). The YJM789 wildtype strain (MATalpha, *lys2*) and its previously generated *aro1* knockout (MATa, *ARO1*::NAT^R^)(Rong-Mullins, Ravishankar, et al., 2017) were used in serial dilution growth assays only as shown. Rich media containing 1% yeast extract, 2% peptone, and 2% dextrose (YPD) or minimal media containing 0.67% yeast nitrogen base and 2% dextrose (YM) with histidine, leucine, uracil, and methionine supplementation for BY4741 strains were used in the various experiments as indicated. Experiments where the aromatic amino acids were supplemented in excess to YPD media were done to a final concentration of 0.02mg/mL tryptophan, 0.03mg/mL tyrosine, and 0.05mg/mL phenylalanine. The strains used in the *TAT2* overexpression serial dilution assay were created by transforming BY4742 yeast with either pRS315 (*CEN, LEU2*) plasmid (Sikorski & Hieter, 1989) or commercially available *LEU2+* multicopy yeast genomic tiling collection plasmids containing genomic regions from chromosome XV corresponding to the region surrounding the *TAT2* ORF (Jones et al., 2008). Supplementations of MCHM, drugs, or other nutrients were added to YPD or YM media as indicated for each experiment. MCHM for experiments is the same crude MCHM formulation that made up the primary chemical presence in the Elk River spill. This formulation is approximately 89% 4-methylcyclohexane methanol, with other cyclohexanes making up the rest of the material was obtained directly from the manufacturer, Eastman Chemical Copmany (Kingsport, TN, US). All concentrations of MCHM indicated in figures are in ppm of crude MCHM formulation, so all conclusions are for this formulation, not pure MCHM.

### Growth and viability assays

Growth and viability assays were performed as previously described (Rong-Mullins, Winans, et al., 2017). Plating for growth assays was only done at a maximum of five strains per plate to keep cells in the central portion of the plate, due to noticed position effects on toxicity, likely from evaporation, a known and observed characteristic of MCHM being its volatility (Phetxumphou et al., 2016; Sain et al., 2015). Furthermore, plates were used within a day to minimize any evaporation of MCHM from the media that would affect concentrations. Concentrations are noted in figure legends. When glutathione was used to attempt to rescue sensitive phenotypes, the concentration on plates was 10mM or 100mM, as noted. Viability assays were carried out for 1.5 to 24 hours while kept in log-phase for concentrations indicated in figures.

### Flow cytometry

BY4741 cells were grown to saturation overnight and returned to mid-log phase. Cells were then diluted to a starting OD_600_ of 0.3 in biological triplicate in YPD media containing 550ppm MCHM. Cells were then harvested at the indicated time. Timepoints longer than 90 minutes required dilutions into fresh treatment media two times over the 12-hour experiment to maintain log-phase. The harvesting procedure involved taking 3mL of culture and fixing and preparing cells as previously published (Haase & Reed, 2002). For cell cycle analysis, cell pellets were fixed in 70% ethanol overnight at 4°C. Fixed cells were pelleted and resuspended in 0.5mL of a 2mg/mL RNAse A solution (50mM Tris pH 8.0, 15mM NaCl boiled for 15 minutes and cooled to room temperature) for a 12-hour incubation at 37°C. Cells were then incubated in 0.2mL of a protease solution (5mg/mL pepsin, 4.5μl/mL HCl) for 20 minutes at 37°C. Finally, samples were stored in 0.5mL of 50mM Tris pH 7.5 at 4°C for up to a week before undergoing flow cytometric analysis. Immediately prior to flow cytometry, samples were shortly sonicated at low power on a Branson model SSE-1 Sonifier for six intervals of one-second bursts, and 50μl of each sample was suspended in 1μM SYTOX Green. Samples were analyzed on a BD LSRFortessa using a FITC channel. Approximately 30,000 events were collected for analysis. FCS Express 5.0 software was used to analyze the DNA content of cells using a multi-cycle DNA histogram and incorporated DNA modeling Statistics. Models were compared via Chi-squared results and the SL S0 model selected for determining the portion of the population in S phase for all samples. All replicates and timepoints fell within good or fair confidence for cell cycle modeling and S phase confidence of the model. The model produced the values for the proportions of the cell population falling within the G1, S, and G2/M phases used for analysis.

For measurement of ROS, live cells were pelleted then suspended in 200μl of 50mM DHE in phosphate-buffered saline (PBS). The dyed cultures were incubated at 30°C for 20 minutes and washed with PBS. A positive control sample of BY4741 cells was treated with 25mM H_2_O_2_ for 1.5 hours. The DHE dyed samples were then analyzed within 2 hours of harvesting on a BD LSRFortessa using preset propidium iodide detection defaults. Approximately 30,000 events were collected per sample for downstream analysis. A High ROS subpopulation range was determined by gating events in downstream analysis at a DHE fluorescence level of 350. This value gave all three biological BY4741 wildtype strain replicates treated as the H_2_O_2_ positive controls approximately 95% of their cells falling above this value (94.69% - 96.86% actual percentages of cells). The wildtype and *vma3* cells, both untreated and treated with MCHM, that fell above a fluorescence level of 350 were considered to be part of a High ROS subpopulation. A one-tailed t-test was used to determine significance of increases in subpopulation percentages for each strain.

### Genetic screen

The genetic screen of the BY4742 haploid knockout collection was performed via the phenotyping of serial dilution growth on solid media by adapting a previous screen technique (Bae et al., 2017). In short, frozen 96-well microtiter plates of the knockout collection were thawed and inoculated via pinning into 96-well growth chambers containing YPD in biological triplicate. Chambers were grown at 30°C for 2 days until all wells were saturated. Each well was then serially diluted into 96-well microtiter plates three times at 20-fold concentrations, for three wells at 20-fold, 400-fold, and 8000-fold dilution from saturated. Diluted plates were then pinned onto large YPD solid media plates with or without 300ppm MCHM. Plates were visually scored for growth at 2-3 days, comparing control, and treated plates for each replicate. Knockout strains showing decreased growth of at least one spot, unaccounted for by decreased growth rate in general on control media, were recorded as screen “hits”. There was occasional variability in growth between replicates, so knockouts that did not replicate sensitive in all three trials filtered out.

### RNAseq

The transcriptomic dataset for this study was produced previously and published with full raw data output and a partial analysis (Gallagher et al., 2020). The raw data are available at GSE, accession number GSE129898 (https://www.ncbi.nlm.nih.gov/geo/query/acc.cgi?acc=GSE129898). A partial reanalysis for this paper included comparisons of specifically strains of the BY4741 untreated control and the BY4741 MCHM treated cells. The library prep, transcript quantification with salmon (v0.9.1), bioinformatics pipeline, and differential analysis with DESeq2(1.18.1) were all performed exactly as in the previous study (Gallagher et al., 2020). GO Term analysis was performed with DAVID bioinformatic database as below for this study.

### Comet Assay

DNA damage resulting from MCHM exposure was quantitated using comet tails as previously described (Azevedo et al., 2011; Oliveira & Johansson, 2012). After incubation in MCHM for 30 minutes or hydrogen peroxide for 45 minutes (Heck et al., 2010; Rowe et al., 2008) as a positive control, single cells were solidified in 1.5% low-melting agarose. The cell walls were digested using 80 μl of 2mg/ ml zymolyase to create spheroplasts and immobilized on a glass slide. After cells were then lysed with 10 mM Tris HCl pH 10, 30 mM NaOH, 1 M NaCl, 50 mM EDTA and 0.05% w/v laurylsarcosine, they were rinsed three times with TAE, and nucleic acid was separated electrophoresis through the gel with 14V for 10 minutes. The gels were then neutralized Tris-HCl pH 7.4, and subsequently fixed with 76% and then 96% ethanol solutions to fix the DNA in place. DNA was stained with ethidium bromide for 20 minutes and then rinsed in water. The DNA fragments formed a tail from the rest of the DNA that has remained intact (the head). Images for each sample 17-42 ‘comets’ were captured with a Nikon A1R confocal microscope and measured via pixel intensity and distance (μm) of the DNA migration through the gel using TriTek CometScore 2.0.0.38. Statistical analysis was performed on tail length measurements using a one-way ANOVA and Tukey’s HSD test in base package of R version 3.6.3 (aov and TukeyHSD commands).

### GO Term analysis

The knockout screen gene list and the RNAseq differential analysis were each analyzed further using the GO Term function of the DAVID bioinformatic database at https://david.ncifcrf.gov (Huang et al., 2009a, 2009b). Default settings were used, and no optional steps were included. Gene lists were searched for all three ontologies, though process and function were determined to show the most informative results and included in the main figures. Full results are available in supplemental files 1-3.

### Data and Reagent availability statement

Raw data for transcriptome analysis are available at GSE, accession number GSE129898 (https://www.ncbi.nlm.nih.gov/geo/query/acc.cgi?acc=GSE129898). GO Term analysis results are available in supplemental files 1-3. The full gene lists of genetic screen hits, significantly upregulated genes, and significantly downregulated genes are available as Tables S1-S3. Haploid BY4741 and BY4742 knockout collection strains and YJM789 *aro1* knockout are available by request. The plasmids and/or resulting strains used in the *TAT2* overexpression analysis are available by request. Crude MCHM used as the main reagent in the experiments was a gift from the Eastman Chemical Company, but a limited supply is available upon request.

## Results

### Determination of S. cerevisiae sensitivity to MCHM treatment

*S. cerevisiae* were treated with a range of MCHM levels to determine minimal concentrations that reduced growth in rich media (YPD). The levels were initially titrated, and approximately 400ppm MCHM inhibited the growth of BY4741 yeast (Figure 1A). The concentrations used on yeast in these growth assays (400-600ppm, 2.8-4.2mM) were approximately 100 times greater than the highest levels recorded in the water distribution system of Kanawha County, WV when monitoring began the day after the spill (Whelton et al., 2014). While the growth assay phenotype itself requires a relatively high dosage of the chemical, the clear phenotype of decreased growth of yeast at this concentration is informative to the design of experiments that exploit the many genomic resources of the yeast. Similar chemical assays of toxic chemicals, such as hydrogen peroxide, use a comparable range of doses (0.5-5mM) to exploit the yeast growth phenotype as a measurement for cellular changes, including the original papers establishing the transcriptional programming of the yeast environmental stress response (0.3mM H_2_O_2_)(Azevedo et al., 2011; Gasch et al., 2000).

**Figure 1:**
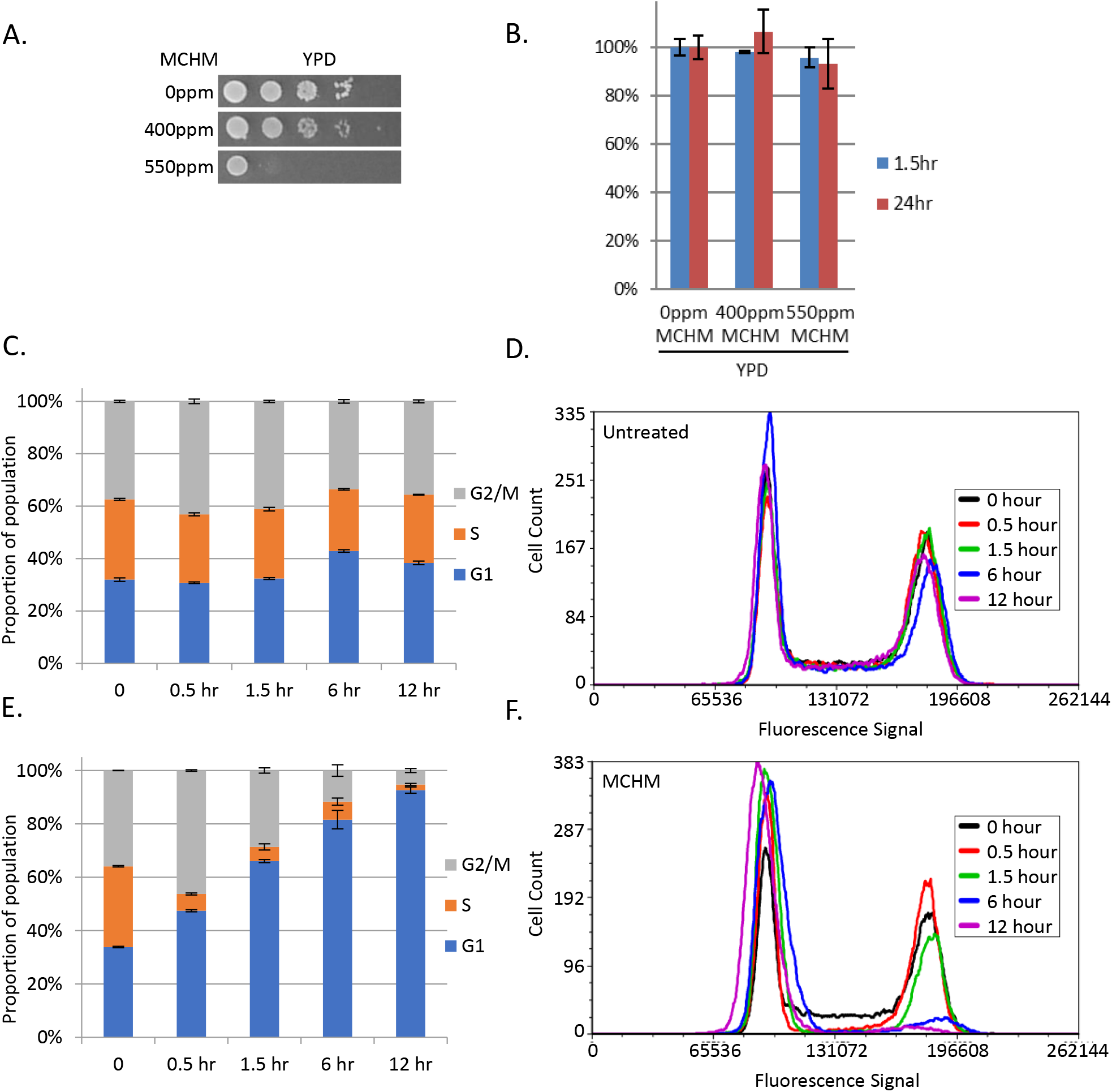
Growth and viability of yeast under MCHM treatment. **A.** Growth of 10-fold dilutions of yeast on YPD media with and without MCHM treatment. BY4741 strain is derived from the S288c background. **B.** Acute viability assay of the MCHM sensitive strain BY4741 treated with MCHM for 1.5 hours and 24 hours. Colony-forming units for three biological replicates were averaged for each treatment then normalized to media without MCHM treatment. In YPD yeast were exposed to 400ppm and 550ppm crude MCHM, and 400ppm and 600ppm grown in YM. P-values are represented by *0.026 **0.018 and ***0.010 **C.** Proportion of log-phase BY4741 in G1, S, or G2/M phases of the cell cycle based on flow cytometric analysis grown in YPD. **D.** Histograms of DNA content in BY4741 population grown in YPD shown in part C, as based on fluorescence intensity of sytox green. **E.** The proportion of log-phase BY4741 in G1, S, or G2/M phases of the cell cycle based on flow cytometric analysis grown in YPD with 550ppm MCHM. **F.** Histograms of DNA content of BY4741 population grown in YPD treated with MCHM shown in part E, as based on fluorescence intensity of sytox green. All samples were analyzed for 0, 0.5, 1.5, 6, and 12-hour time points to monitor cell cycle changes over time.

### Decreased growth of yeast treated with MCHM is not due to large-scale cell death

To assess whether MCHM inhibited growth by decreasing cellular viability or by arresting growth, acute and chronic viability assays were completed (Figure 1B). Mid-log phase yeast were acutely exposed for 90 min and chronically exposed for 24 hours to the 550ppm MCHM. Yeast were then washed and plated onto YPD with no MCHM. After growing plates for two days, there was little if any reduction in colony-forming units with the YPD-MCHM treated yeast.

To determine if the reduction in growth is due to cell cycle arrest, we analyzed MCHM treated yeast by flow cytometry (Figure 1C-1F). Untreated cellular populations showed an asynchronous pattern, for which each of the three phases had upwards of 20% of the total cellular population (Figure 1C and 1D). However, within 30 minutes, the MCHM treated samples showed that less than 10% of the population was in S phase (Figure 1E and 1F). The cells treated with MCHM appear to arrest almost immediately in G1 phase, leading to a nearly instantaneous loss of S phase population as no new cells leave G1. Over the course of the 12-hour experiment, the population in G2 also decreased to less than 10% of cells (Figure 1E and 1F). MCHM treated yeast failed to grow because they arrested in G1, not because of decreased viability.

### Genetic screen reveals cellular pathways and components required for MCHM tolerance

The cell cycle arrest of MCHM treated yeast was hypothesized to be attributable to environmental stress response programming, as many stressors initiate arrest as the cells attempt to ameliorate and recover from damage caused (Gasch et al., 2000). In order to test this hypothesis, we collected genomic datasets to analyze which genes were functionally required for MCHM tolerance in mutant strains and which were differentially expressed in wildtype cells. The nearly 5000 strains of the haploid BY4742 knockout collection were tested for growth phenotypes on 300ppm MCHM in a genetic screen (Supplemental Figure 1). The results revealed that 329 genes were required for the tolerance to MCHM treatment (Supplemental Table 1). Several individually important environmental stress response genes were revealed by the screen. Two genes of note for their role in the control of Msn2/4-regulated stress responses were *MCK1* and *MSN5*. Deletions of these two genes have been shown previously to affect the ability of Msn2 to control stress response programming (Gutin et al., 2019).

GO term analysis of the results of the screen pointed to mitochondrial translation as the most enriched subset of yeast genes (Figure 2A). This particular GO term agrees well with previously published data showing that petite yeast strains are sensitive to MCHM (Pupo, Ayers, et al., 2019). Mitochondrial function is important for dealing with many stressors, especially those involved in reactive oxygen species production. The next most enriched gene subset was the components of the vacuolar ATPase (v-ATPase). This transporter maintains the acidity of the vacuole by pumping H+ ions across the vacuolar membrane. The homeostatic processes that use this ion gradient are evidence that MCHM may be causing ROS stress(N. Nelson & Harvey, 1999; Nishi & Forgac, 2002).

**Figure 2:**
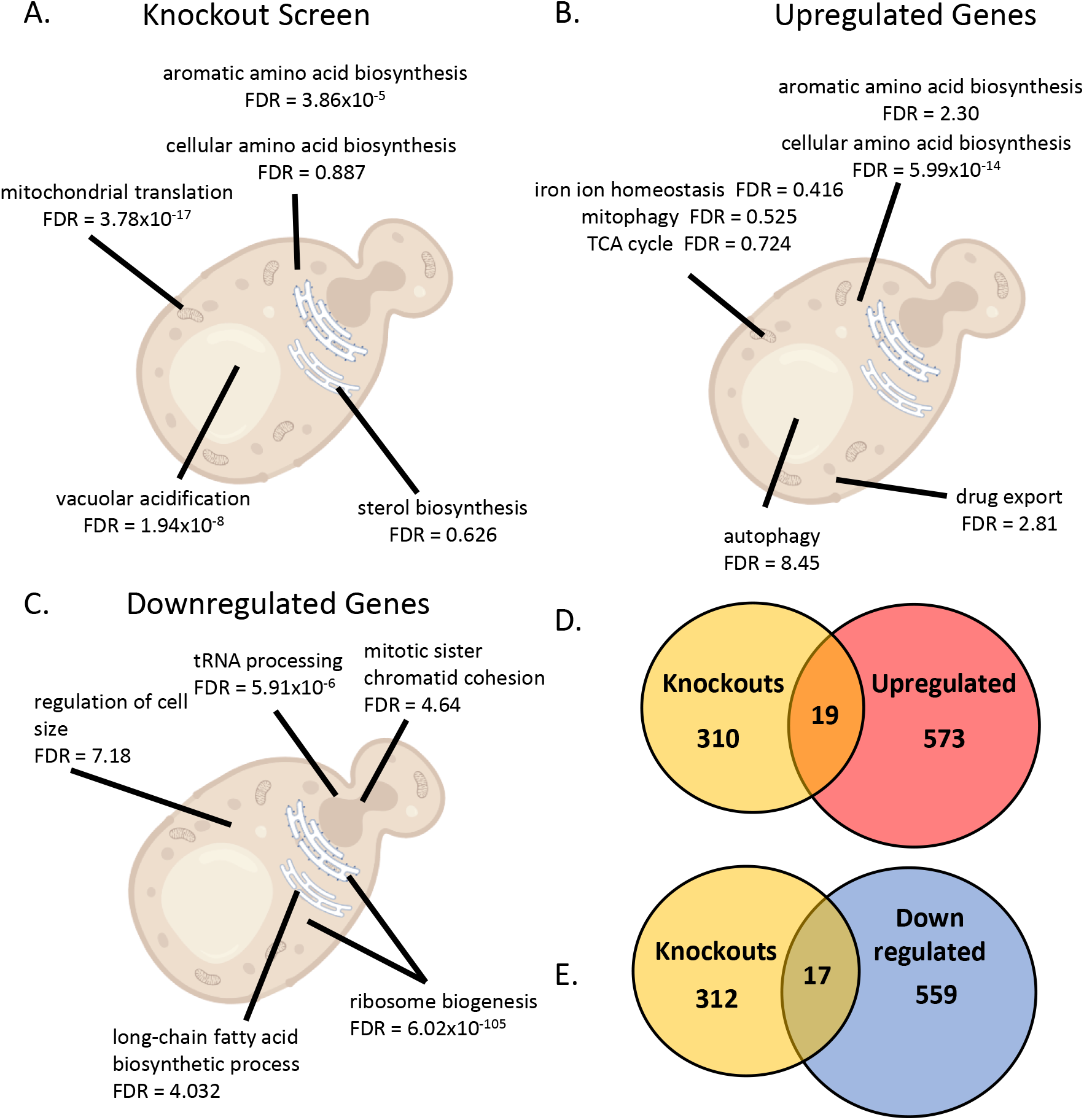
Genetic screen of knockout collection and RNAseq of BY4741 gene lists. **A.** Select top biological process GO Terms for the list of 329 genes found to be hits in the genetic screen of BY4742 knockout collection. FDR is shown for each term. **B.** Select top biological process GO Terms for the 592 significantly upregulated genes from BY4741 treated with MCHM versus untreated YPD. FDR is shown for each term. **C.** Select top biological process GO Terms for the 572 significantly downregulated genes from BY4741 treated with MCHM versus untreated YPD. FDR is shown for each term. **D.** Venn diagram of the overlap between the 329 genes from the genetic screen and the upregulated genes. **E.** Venn diagram of the overlap between the 329 genes from the genetic screen and the downregulated genes.

The next most enriched GO terms were for amino acid biosynthesis, especially aromatic amino acid biosynthesis. As the screen was performed on YPD media, it was surprising that these biosynthetic genes, including early precursor producing enzymes (*ARO1* and *ARO3*) and nearly the entirety of the tryptophan biosynthetic pathway (*TRP2-5*), were required when presumably excess amino acids were available from the media.

### MCHM treatment significantly affects the transcription of large portions of the yeast genome

Transcriptomic analysis of the BY4741 strain treated with MCHM also supported the hypothesis that the environmental stress response was activated with exposure to MCHM. There were 592 significantly upregulated genes in response to MCHM (Supplemental Table 2 and Figure 2B), while there were 576 genes significantly downregulated (Supplemental Table 3 and Figure 2C). These lists of genes were analyzed for GO terms and several of the terms were consistent with the activation of the ESR (Figure 2B-C). The upregulated GO terms for drug export, autophagy, and mitophagy point to ESR mechanisms for the removal of toxic substances from the cell and the reapportionment of cellular resources to overcome and adapt to a stressor. The upregulated terms related to amino acid biosynthesis and mitochondrial function (TCA cycle, iron ion homeostasis) were consistent with the genetic screen GO term analysis. The mitochondrial overlap between the datasets is likely due to the importance of ROS stress response, though the importance of the amino acid pathways as mentioned above, especially considering the cells are growing in rich media, is unclear.

The clearest indication for the central role of ESR activation by MCHM exposure was the downregulated GO terms. Ribosome biogenesis and related functions (other ribosomal terms were also significant as seen in Supplemental Table 3 but omitted from the figure for legibility) were the primary GO terms found in this dataset. They are also classic pathways downregulated in the ESR (Gasch et al., 2000). Most ribosomal biogenesis protein genes are essential and do not appear in the haploid knockout collection used in the genetic screen, with 81% of the 79 ribosomal protein genes essential for growth in standard conditions (Steffen et al., 2012). Therefore, the transcriptomic data was necessary supporting evidence for ESR involvement in MCHM response in combination with the stress gene and amino acid biosynthesis knockouts appearing in the screen. Other terms from the downregulated genes, including fatty acid biosynthesis and regulation of cell size, are also likely related to the observed cell cycle arrest and the reapportionment of energy and resources required for overcoming stressors.

### Combined genomic datasets implicate nutrient starvation

The genetic screen and transcriptome datasets were compared to identify shared genes that might be especially important to MCHM response. The list of overlapping genes from the screen and the upregulated datasets included *SNQ2*, an ABC multidrug transporter associated with the removal of foreign substances from the cell (Figure 2D) (Decottignies et al., 1995; Decottignies & Goffeau, 1997; Rogers et al., 2001; Servos et al., 1993). While there are several such transporters (and related transcription factors) found in the transcriptomics, this is the only one found in the screen to be required for the tolerance of MCHM. We hypothesize that this transporter is responsible for the removal of MCHM or a related toxic byproduct from the cell.

Other upregulated genes also found in the knockout screen are involved in mitochondrial function and ROS stress responses. *GGC1* and *TGL2* encode mitochondrial proteins involved in GTP/GDP transport and triacylglycerol lipase activity at the mitochondria respectively(Ham et al., 2010; Vozza et al., 2004). The *GGC1* knockout in particular produces petite yeast which are known to be sensitive to MCHM (Pupo, Ayers, et al., 2019; Vozza et al., 2004). The *POS5* gene, identified originally for conferring peroxide sensitivity, is a mitochondrial protein that acts as an NADH kinase(Krems et al., 1995; Strand et al., 2003). It is required for maintenance of mitochondrial DNA, the loss of which results in the aforementioned petite yeast, as well as detoxification of ROS in the mitochondria(Strand et al., 2003). The presence of these genes in both datasets points to the importance of mitochondrial function in the response to MCHM stress, most likely due to ROS production.

The other major insight of the combined screen and upregulation datasets is the presence of several amino acid biosynthetic genes. Many of these genes fall under the control of the general amino acid activator Gcn4 (Braus, 1991). Several single genes from pathways for isoleucine/valine (*ILV6*), methionine (*MET22*), and lysine (*LYS2*) were found in the overlap. Furthermore, the aromatic amino acid biosynthetic genes *ARO1* and *ARO3*, as well as four out of five tryptophan biosynthetic genes (*TRP2, TRP3, TRP4,* and *TRP5*), were found in both datasets. As tryptophan pathways in particular, but also tyrosine, have previously been implicated in the response to multiple types of stress including nutrient starvation, pH, metal ions, SDS, and DNA damaging chemicals, it was initially unclear which effect MCHM might be having on the cell to result in the sensitivity of these mutants (Godin et al., 2016; González et al., 2008; Schroeder & Ikui, 2019).

The downregulated dataset overlaps with the screen revealed a few genes with a possible relationship to the transcriptional activator Gcn4 (Figure 2E). *BUD27* is related to translation initiation and may be involved in the expression of genes controlled by Gcn4 (Deplazes et al., 2009; Gstaiger et al., 2003; Mirón-García et al., 2013). *GUA1* is negatively regulated by nutrient starvation and its mutants have been shown to be impaired in *GCN4* translation (Escobar-Henriques et al., 2003; Escobar-Henriques & Daignan-Fornier, 2001; Iglesias-Gato et al., 2011). *PRM7* also contains Gcn4 binding elements, connecting it to control by this transcriptional activator (Schuldiner et al., 1998). While not directly under Gcn4 control, *PRS1*, *PRS3*, and *TAT1* provide more evidence for the significance of tryptophan and amino acid metabolism in MCHM sensitivity. *PRS1* and *PRS3* encode 5-phospho-ribosyl-1(alpha)-pyrophosphate (PRPP) synthetase genes which are necessary for the production of purines and pyrimidines, and histidine and tryptophan amino acids (A. T. Carter et al., 1997; Andrew T. Carter et al., 1994; Hernando et al., 1999). While there are five paralog PRPP synthetases in yeast (Prs1-5), different multimeric complexes of the paralogous proteins are likely involved in producing wildtype levels of PRPP. Disruptions in *PRS1* and *PRS3* have the largest reduction in PRPP pools for nucleotide and amino acid production (Hernando et al., 1999). The protein product of the *TAT1* gene is an amino acid transporter, characterized originally for its high-affinity import of tyrosine, but also shown to be a low-affinity transporter of tryptophan (Schmidt et al., 1994). The combined screen and transcriptomic profiles indicated that MCHM may cause yeast cells to undergo nutrient starvation, possibly nitrogen specifically, despite growing in rich media. Cells may be responding by initiating cell cycle arrest and reapportioning nitrogen resources until recovered.

The importance of the aromatic amino acid synthetic genes in MCHM tolerance was investigated by additional supplementation of tryptophan, tyrosine, and phenylalanine in rich media containing MCHM. This was insufficient to rescue the growth of either wildtype or pathway gene knockout strains in MCHM (Figure 3A). It is possible that the excess tryptophan was not sufficiently imported into the cells based on the sorting of transporters such as Tat2 to the vacuole instead of the membrane (Schroeder & Ikui, 2019). This is plausible as ergosterol levels in the cell affect this sorting preference, and MCHM is known to alter sterol levels in the cell (Pupo, Ku, et al., 2019; Umebayashi & Nakano, 2003). Overexpression of *TAT2* using a multicopy tiling plasmid did not rescue growth on MCHM with or without supplemental aromatic amino acids (Supplemental Figure 2). The levels of several amino acids increase with MCHM treatment, including the levels of tyrosine (Pupo, Ku, et al., 2019). Unfortunately, the levels of tryptophan were not measured by GC-MS, so it is unknown if levels change with MCHM treatment. Yeast cells seem to be experiencing a nutrient starvation signal upon exposure, reacting by increasing production of amino acids in a way that is required according to knockouts from the screen, and yet failing to adapt to the chemical, based on the presence of excess amino acids alone.

**Figure 3:**
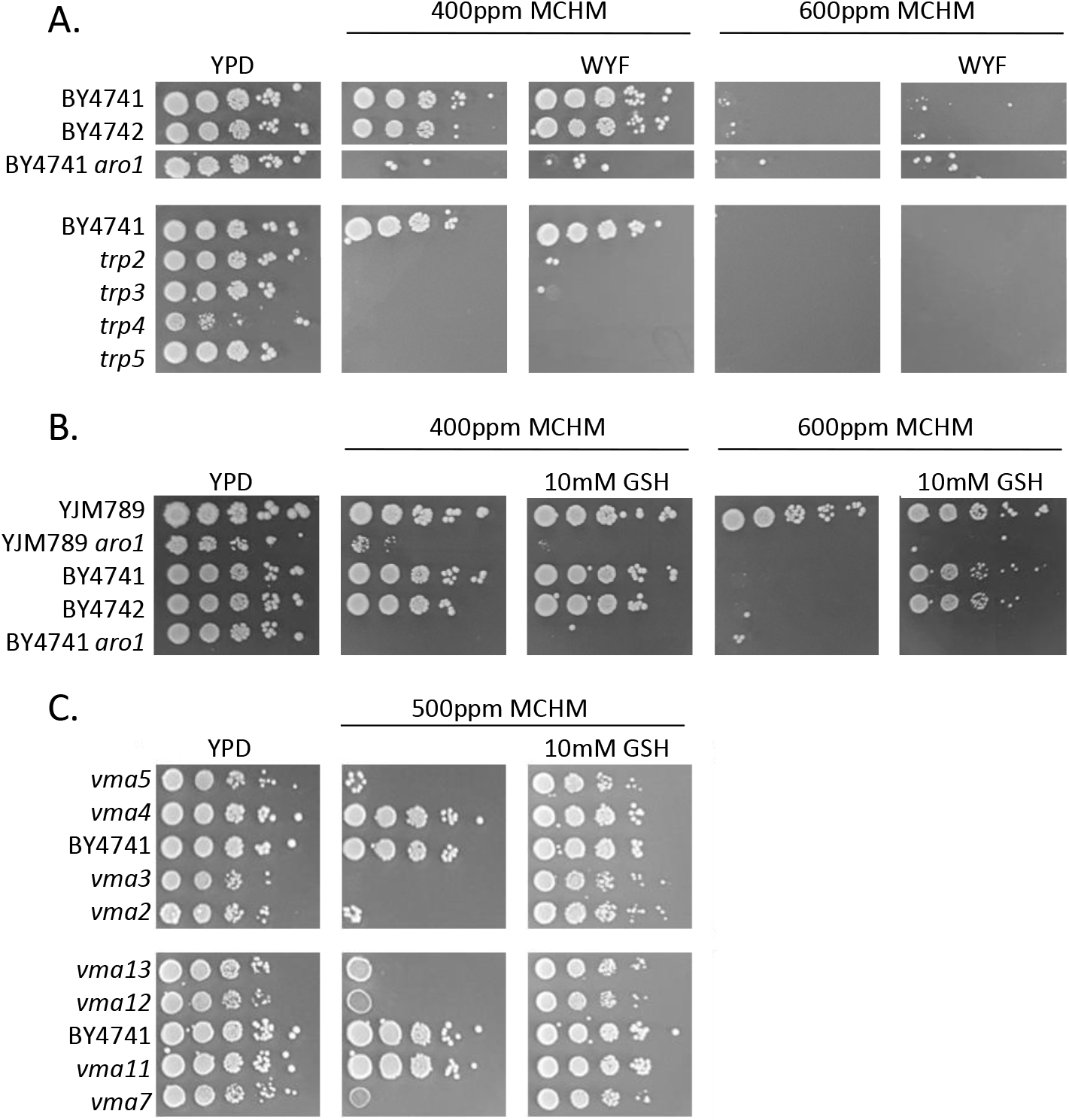
Sensitive mutants from gene lists cotreated with possible rescuing chemicals. **A.** 10-fold serial dilution growth assay of knockouts of aromatic amino acid biosynthesis genes found in the genetic screen. Strains were cotreated with MCHM and aromatic amino acids Tryptophan (W), Tyrosine (Y), and Phenylalanine (F) in rich media, testing the extra supplementation of these amino acids beyond normal levels in YPD. **B.** 10-fold serial dilution growth assay of wildtype strains and knockout of aromatic amino acid biosynthetic gene *ARO1* with and without cotreatment of 10mM antioxidant glutathione (GSH) and MCHM. **C.** 10-fold serial dilution growth assay of knockouts in the vacuolar acidification pathway as found in the genetic screen. Strains were cotreated with and without 10mM antioxidant GSH and MCHM.

### Uncharacterized ORFs both upregulated by and required for MCHM responses

Of note in the combined datasets is the requirement of *YBR241c* for tolerance to MCHM (Table 1). Ybr241c is a paralog of the vacuolar sorting protein, Vps73, and is localized to the vacuolar membrane (Matsumoto et al., 2013). Acidification of the vacuole is critical for metabolizing chemicals such as MCHM and Ybr241c relocalizes to the cytoplasm when acidification of the vacuole is blocked (Matsumoto et al., 2013). Ybr241c also physically interacts with Gtt1, a glutathione transferase of vacuolar proteins (Chandel et al., 2016; Yu et al., 2008). Like Vps73, Ybr241c shares homology with other sugar transporters such as Hxt1-17 and related proteins. As Ybr241 is a <u>v</u>acuolar protein <u>v</u>ital for <u>s</u>tress response, we named it Vvs1. Ycr061w localizes to cytoplasmic puncta that do not appear to be vacuoles but instead vesicles (Huh et al., 2003; Tkach et al., 2012). Ycr061w contains 10-11 predicted transmembrane domains (Weill et al., 2019). As Ycr061w is a transmembrane protein vital for stress response, we named it Tvs1.

**Table 1:** List of genes overlapping between the genetic screen and upregulated or downregulated RNAseq results, as shown in Figure 2. Genes are organized, with subheadings, to highlight functions relevant to environmental stress responses.

| Overlap of Screen and Upregulated Genes | Overlap of Screen and Downregulated Genes |
| --- | --- |
| Aromatic Amino Acid Biosynthesis: | Aromatic Amino Acid Synthesis/Transport: |
| *ARO1* | *PRS1* |
| *ARO3* | *PRS3* |
| *TRP2* | *TAT1* |
| *TRP3* |  |
| *TRP4* | RNA Processing/Ribosome Biogenesis: |
| *TRP5* | *MRT4* |
|  | *NRS1* |
| Amino Acid Biosynthesis: | *RNR1* |
| *ILV6* | *RPA1* |
| *LYS2* | *TSR2* |
| *MET22* |  |
|  | Translation: |
| Mitochondrial Localization: | *BUD27* |
| *GGC1* | *FUN12* |
| *POS5* |  |
| *TGL2* | Protein Chaperone: |
|  | *YDJ1* |
| ABC Multi-drug Transporter: |  |
| *SNQ2* | Transcriptional Regulator of Stress Response: |
|  | *MOT3* |
| Transcription of rDNA: |  |
| *RRN10* | Ion Transporter/Metal Homeostasis: |
|  | *PHO84* |
| Miscellaneous Function: |  |
| *MOG1* | Miscellaneous Function: |
| *XYL2* | *BUD30* |
| *VVS1 (YBR241C)* | *GUA1* |
| *TVS1 (YCR061W)* | *PRM7* |
| *YHI9* | *SAM1* |

### Reactive Oxygen Species production is implicated by growth rescue via glutathione

As mitochondrial processes and iron ion homeostasis were found to be significant results in the genomic datasets, we further explored the role of reactive oxygen species in MCHM-induced stress. Glutathione serves as an antioxidant for the cell, particularly in the mitochondria and ER, and is also important in iron-sulfur homeostasis (Bulteau et al., 2012). It was hypothesized that treatment with glutathione may rescue ROS stress created by exposure to MCHM, so yeast were co-treated with the chemicals. Glutathione was sufficient to rescue the growth of wildtype yeast, even at extremely high doses of 600ppm (Figure 3B). Tryptophan levels are important for resistance to DNA damaging agents (Godin et al., 2016). However, glutathione was not able to rescue the sensitivity of the *aro1* mutant, so the unknown non-amino acid role in MCHM tolerance of the aromatic amino acid biosynthetic genes does not seem to be related to an ROS stress mechanism. The sensitivity of the *aro1* knockout mutant was replicated in a divergent yeast strain background, YJM789 (Figure 3B), to control for genetic background specific sensitivity.

The yeast vacuolar ATPase was the second most significant GO term from the genetic screen dataset. The v-ATPase is responsible for acidifying the vacuole of yeast cells by hydrolyzing ATP in order to pump hydrogen ions into the vacuole, creating an electrochemical gradient with multiple homeostatic roles (Deranieh et al., 2015; H. Nelson & Nelson, 1990; N. Nelson & Harvey, 1999; Nishi & Forgac, 2002; Ohya et al., 1991). *vma* mutants show at least three sensitivity phenotypes that may be related to MCHM’s effects, including metal ion homeostasis, pH sensitivity, and oxidative stress sensitivity. To test if oxidative stress may be responsible for the sensitivity of *vma* mutants, we co-treated these mutants with MCHM and glutathione (Figure 3C). The observed growth rescue with glutathione treatment supports the hypothesis that the v-ATPase is required to provide a robust level of ROS protection yeast cells need to survive MCHM.

### Biochemical assays reveal direct evidence for the presence of ROS in MCHM treated cells and point to mutant sensitivity as reliant on ROS homeostatic capacity

As genomic datasets and mutant growth assays had implicated ROS production as an effect of MCHM treatment, we attempted to directly detect ROS inside the cell. The dye dihydroethidium (DHE) reacts with intracellular ROS and then creates a fluorescent signal. We dyed yeast treated with MCHM with DHE, then detected the changes in fluorescent signal in the populations via flow cytometry. The flow cytometry results revealed an increase in fluorescence for a subpopulation of wildtype cells (Figure 4A). Within the genetic clonal population of MCHM treated cells represented by the red curve, phenotypic heterogeneity existed. The majority of MCHM treated cells did not have an increase in ROS levels, in contrast to hydrogen peroxide treated yeast (gray line). However, a subpopulation of the red curve, as seen in the portion of the curve under the marker labeled High ROS, did have increased ROS. The High ROS subpopulation of cells increased from 2.96% to 7.82% of the total cellular population in the BY4741 strain (Supplemental Figure 3). It is a known phenomenon that clonal populations in liquid culture can show phenotypic heterogeneity, specifically with respect to ROS sensitivity(Sumner et al., 2003). This may explain why there is little reduction in the viability of BY4741 cells seen in Figure 1B. If the cells that successfully arrest their cell cycle were able to adapt and minimize their ROS production to avoid MCHM-induced death, they may appear as the large subpopulation with a lower ROS signal in the data. The cells that failed to do so may represent both the few cells that die in the viability assay, as well as those appearing as the small peak in the High ROS range. This assay revealed the first direct evidence that MCHM treatment produces ROS in the cell, in agreement with the indirect evidence originally seen in the sensitivity of *vma* mutants and glutathione rescue.

**Figure 4:**
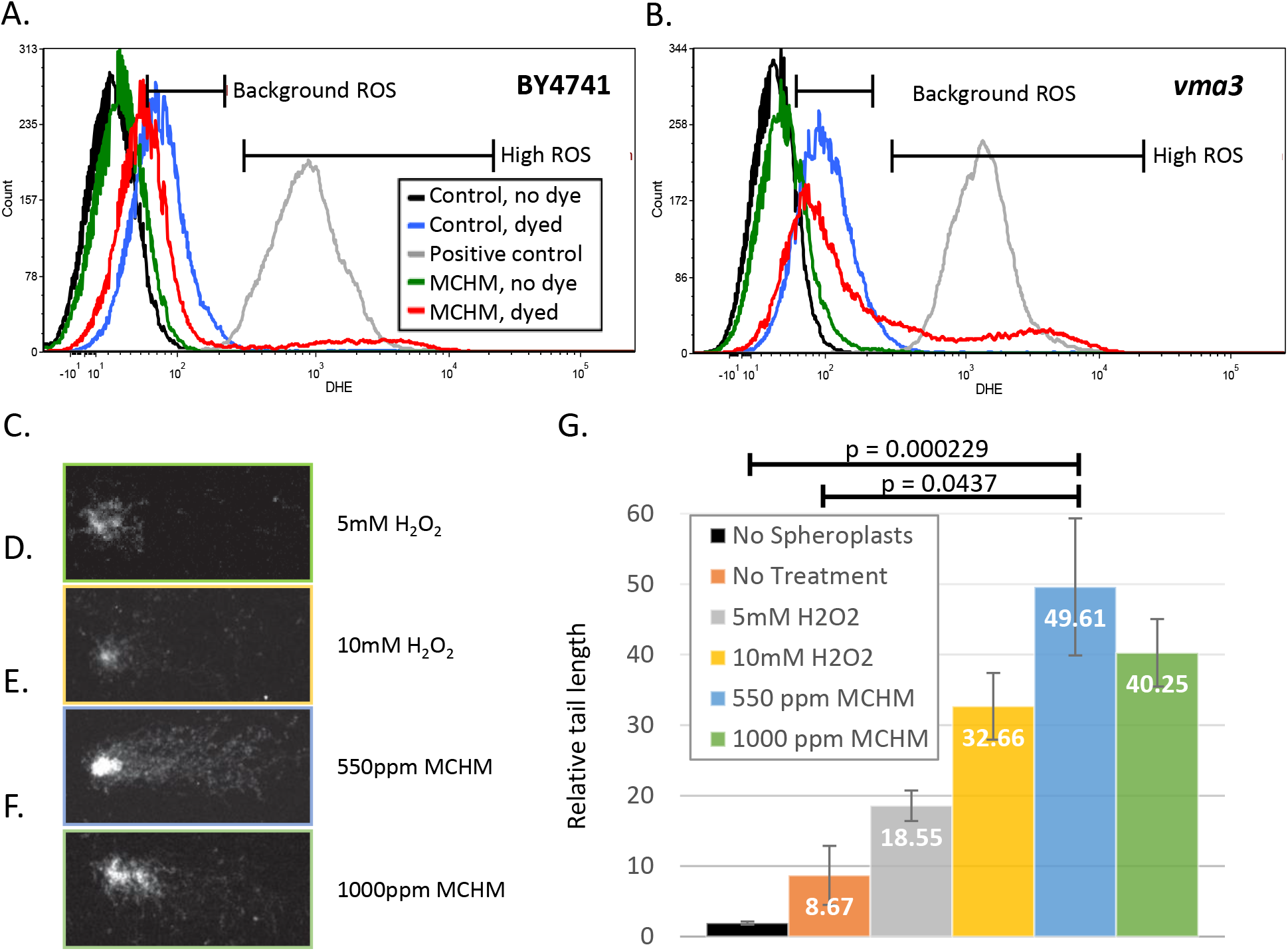
Flow cytometry and comet assay of yeast strains reveals the production of Reactive Oxygen Species and DNA damage. Levels of ROS in strains of yeast exposed to MCHM, based on fluorescence of the ROS-reactive dye DHE. Yeast were incubated for 12 hours with or without MCHM then stained with DHE for 20 minutes before cells were sorted using flow cytometry. Hydrogen peroxide treatment for 1.5 hours was used as a positive control to generate ROS in control cells (grey line). Background ROS of untreated yeast in blue measures endogenous ROS compared to unstained yeast with no DHE (black). MCHM treated yeast are in green while MCHM treated yeast stained with DHE are in red. Markers were inserted on the histogram to indicate a range of fluorescence corresponding to background or high ROS based on peaks for untreated control cells. Histograms are for the strains **A.** BY4741, and **B.***vma3* knockout mutant. **C-F.** Examples of cells visualized by microscopy for the comet assay. Cells underwent treatment prior to spheroplasting, electrophoresis on a slide, and visualization via microscopy and ethidium bromide staining. Lengths of tails were measured with TriTek CometScore 2.0.0.38 software. The samples shown are examples of the following treatments **C.** 5mM H_2_O_2_, **D.** 10mM H_2_O_2_, **E.** 550ppm MCHM, and **F.** 1000ppm MCHM. **G.** Graph of tail lengths of cells determined via comet assay.

While the effects of MCHM on yeast point to an increase in ROS stress for the cell, wildtype cells seem to be robust enough, on average, to limit this stress, while mutants in certain pathways cannot. To test this hypothesis, we also performed the DHE assay on one of the most sensitive knockout mutant strains, *vma3*, which previously showed rescued growth with glutathione treatment. This mutant strain showed a similar pattern of DHE fluorescence when treated with MCHM, with the appearance of a High ROS population peak (Figure 4B). The major difference from the wildtype was an increase in the size of the High ROS population peak. The High ROS population in the *vma3* strain rose from 1.08% to 22.19% of all cells (Supplemental Figure 3). The mutant strain also appeared to show a right-shift of the background ROS populations as well. This may indicate *vma* mutants have higher ROS levels in unstressed conditions. Therefore, these strains have an innate sensitivity to MCHM due to their inability to maintain ROS homeostasis. We hypothesize that robust cells were able to detect the increased ROS caused by MCHM stress, then adapt to this by arresting growth and limiting their background ROS production.

### MCHM treatment causes DNA damage to wildtype yeast cells

We also questioned whether the ROS produced in MCHM treated cells may cause DNA damage, a known complication in ROS stressed cells (reviewed in Ayer et al., 2014; Brennan et al., 1994; Lafleur & Retèl, 1993; Perrone et al., 2008; Temple et al., 2005). To address this question, a comet assay was performed to directly measure damage. This assay uses microscopy to measure the length of “tails”, resembling comets, released from spheroplasted and lysed cells, and migrated across agarose covered slides via electrophoresis (Figure 4C-F). Longer tails indicate DNA fragments, presumably from damage resulting in double-stranded breaks. Yeast cells treated with MCHM concentrations of 550 ppm showed significantly longer comet tails as compared to no treatment and no spheroplast controls (Tukey’s test p adj = 0.0437 and 0.000229 respectively) and approached significance as compared to 5mM H_2_O_2_ positive controls (p adj = 0.0655). This is the first direct evidence of DNA damage from MCHM treatment, confirming implications of data such as the ROS assay, cell cycle arrest, and other stress response data in previous studies (Lan et al., 2015).

## Discussion

The evidence presented here supports a model where MCHM activates the general environmental stress response programming of the yeast cell via Msn2/4 transcription factors, including specific responses consistent with ROS stress and nutrient starvation. Cell cycle arrest begins within 30 minutes of treatment, likely due to the initial signaling of nutrient starvation. These signals inactivate growth mechanisms soon after the cell is exposed, but before ROS accumulate. The ROS stress accumulates over six to twelve hours and the asynchronous population of cells continues to arrest in G1 during this time period. When able to mount a robust response, as in wildtype strains, cells were able to manage the stress, recover, and return to proliferation (Figure 5) at dosage levels significantly higher than those found in the Elk River spill. In this model, several signals initiate multiple stress responses, including nutrient starvation, reactive oxygen species, and DNA damage itself.

**Figure 5:**
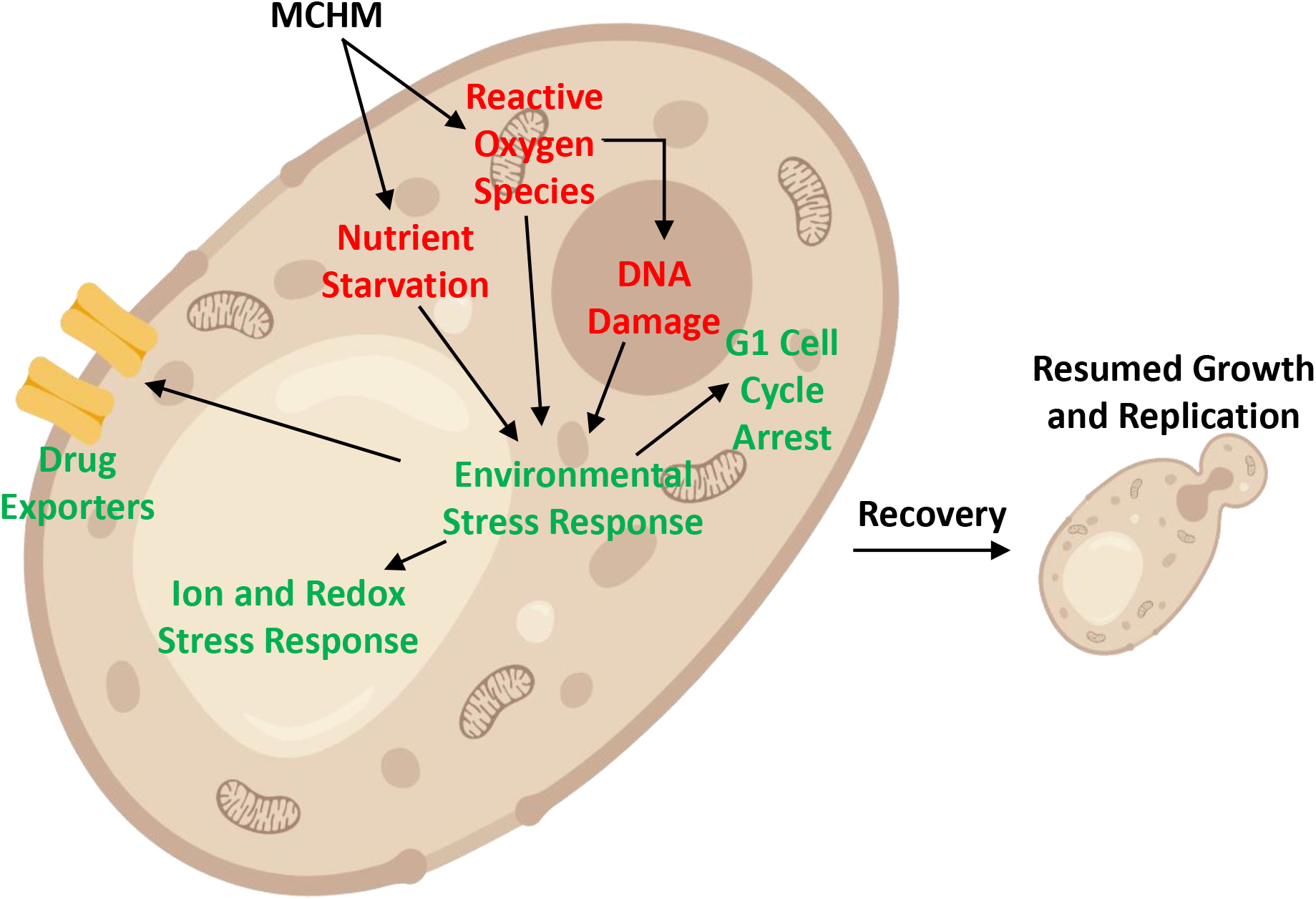
Model of response and tolerance of yeast cells to MCHM treatment. Diagram of yeast cell showing MCHM entering cell and causing ROS, DNA damage, and possible nutrient starvation. These signals are shown triggering environmental stress response leading to cellular responses that lead to recovery and resumption of replication.

There is direct evidence of ROS production, DNA damage, and rescue by antioxidant treatment. Other data have previously indicated that petite yeast, known to be sensitive to ROS, are also sensitive to MCHM, and this agrees with the sensitivity of the mitochondrial translation mutants revealed in the genetic screen data of this study (Pupo, Ayers, et al., 2019). The vacuolar ATPase mutants, which are sensitive to metal and pH stress, as well as ROS, were one of the other most enriched GO terms from the screen. This study, combined with previous data, points to MCHM as a source of perturbation for metal ions and ROS (Lan et al., 2015; Pupo, Ayers, et al., 2019).

Evidence of nutrient starvation is less clear, as previous work reveals that several amino acid levels increase in the cell (Pupo, Ku, et al., 2019), and treatment with excess aromatic amino acids cannot rescue growth. However, the cell’s programmed response to nutrient starvation is initiated by the cell to respond to MCHM, as wildtype cells upregulate the cellular and aromatic amino acid biosynthetic pathways. Furthermore, mutants lacking genes in these pathways, those in aromatic amino acids and tryptophan, in particular, are more susceptible to MCHM than wildtype cells, even in rich media with excess aromatic amino acids available. We are able to eliminate the possibility that there is a ROS homeostatic role that these aromatic amino acid biosynthesis mutants are unable to perform, as treatment with glutathione was unable to rescue their sensitivity, unlike mutants of the vacuolar ATPase.

The major question remains why the aromatic amino acid biosynthesis genes, including all but one of the tryptophan pathway genes, are required for resistance to the chemical. Supplementation with excess tryptophan in YPD did not rescue growth on MCHM, but it is not clear if this supplementation increased intracellular levels of tryptophan. It is possible the sorting of the transporter Tat2 to the plasma membrane was insufficient to import excess tryptophan. Furthermore, GC-MS failed to measure intracellular levels of tryptophan. While overexpression of *TAT2* was previously sufficient to rescue SDS sensitivity in *mck1* knockout mutant cells(Schroeder & Ikui, 2019), a multicopy plasmid expressing the *TAT2* region of chromosome XV was not able to rescue growth on MCHM. Another hypothesis is that the biosynthetic genes themselves are important for signaling recovery from stress so that cells may resume proliferation, regardless of the actual levels of aromatic amino acid products related to the pathway. The *trp1-5* mutants are sensitive to rapamycin (González et al., 2008), so it is possibly a recovery phenotype. The chemical glutathione can rescue MCHM treatment in wildtype cells, but it cannot do so in these mutants. Therefore, these genes may be required to signal the cells are recovered and ready to proliferate following the glutathione treatment amelioration of ROS stress. Another possibility is that there is a separate source of stress besides nutrient availability or ROS that the genes are required to address, such as production or conversion of other small molecules. It would be useful to determine if there are other stressors for which these genes are required, regardless of amino acid levels, to address the possibility that this is specific to MCHM exposure.

The Elk River MCHM spill revealed that there are many things we do not know about the industrial chemicals that we live among. This and other studies since have revealed that levels of the chemical may not be acutely toxic, but there are immense and still only partially characterized effects on the biochemical pathways that control eukaryotic metabolism. This chemical causes the production of reactive oxygen species and DNA damage in yeast and alters the transcriptome and metabolome in extensive ways. While the effects on possible nutrient starvation and associated amino acid biosynthesis are not fully understood, cells with robust homeostatic processes are generally able to activate the environmental stress responses programmed in their genes to recover from MCHM exposure and resume proliferation.

## Acknowledgments

The MCHM sample used was obtained as a gift from Eastman Chemical Company. The yeast knockout collection was a gift from Angela Lee. Yeast cartoon and diagram images in Figures 2A, 2B, 2C, 5, and Supplemental Figure 1 were created with BioRender.com. We would like to thank the WVU Flow Cytometry Core, supported by the following equipment grants: S10OD016165, P30GM103488, and P20GM103434. We would like to thank the WVU Microscope Imaging Facility for use of the Nikon A1R confocal microscope. The imaging facility is supported by the following WVU Cancer Institute and NIH grants: P20RR016440, P30GM103488 and U54GM104942, and the Nikon confocal microscope is supported by U54GM104942 and P20GM103434. We would like to acknowledge the WVU Genomics Core Facility, Morgantown WV for the support provided to help make this publication possible. We would like to acknowledge the WVU Genomics Core Facility, Morgantown WV for the support provided to help make this publication possible and CTSI Grant #U54 GM104942 which in turn provides financial support to the Core Facility. Amaury Pupo analyzed and mapped raw transcriptome reads and analyzed differential expression. This work was supported by NIH NIEHS R15ES026811-01A1. MCA was supported by a WVU STEM Mountains of Excellence Fellowship.

## Author Contributions

MCA designed and carried out yeast experiments and wrote the paper. ZNS performed comet assay. JEGG designed the study, supervised the project, and co-wrote the paper.

## Figure Legends

**Supplemental Figure 1: *Diagram of the genetic screen of haploid BY4742 knockout collection.*** The 96-well plates of the knockout collection were each inoculated into growth chambers in triplicate by pinning and grown for two days to saturation. These chambers were then serially diluted 20-fold three times into new 96 well plates, before being pinned onto YPD agar plates with or without 300ppm MCHM. Pictures were taken after 3 days of growth and scored for growth defects consisting of decreased growth of at least a full spot on MCHM plates compared to control strain (wt BY4742) and control plate growth.

**Supplemental Figure 2: *TAT2 overexpression does not rescue the MCHM growth defect.*** 10-fold serial dilution growth assay of BY4742 with indicated plasmids on minimal and rich media with indicated treatment with MCHM and supplemental aromatic amino acids tryptophan (W), tyrosine (Y), and phenylalanine (F). pRS315 is a control CEN plasmid expressing the *LEU2* marker to confer growth on leu-media to the wild-type BY4742 without overexpression of any genomic regions. *TAT2* overex. contains a tiling array multicopy plasmid with the genomic region of chromosome XV containing *TAT2* for its overexpression (from the *IFM1* ORF to the left of *TAT2* to the YOL019W-A ORF to the right, six complete ORFs and one partial). The control region right plasmid is the same multicopy tiling array plasmid but with the genomic region flanking the *TAT2* ORF on the right side, from within the *TAT2* ORF to the YOL015W ORF (six complete ORFs and two partials). The control region left plasmid is the same multicopy tiling array plasmid but with the genomic region flanking the *TAT2* ORF on the left side, from the *IFM1* ORF to the *DIS3* ORF directly to the left of *TAT2* (two complete ORFs and two partials).

**Supplemental Figure 3: *Increased percentage of High ROS cells with MCHM treatment.*** Percentages of High ROS subpopulation of cells from the BY4741 wiildtype and *vma3* mutant strains, based on fluorescence of the ROS-reactive dye DHE are shown. Yeast were incubated for 12 hours with or without MCHM then stained with DHE for 20 minutes before cells were sorted using flow cytometry. Fluorescence level (350) for determining High ROS subpopulation was based on approximately 95% (94.69%-96.86%) of H_2_O_2_ positive control cells in all three BY4741 replicates being above this fluorescence level. Bar graph shows the averages of three biological replicates for each strain and treatment with standard error bars. One-tailed t-test was used to determine p-values.

**Table S1: *Full list of knockout strains found as hits in the genetic screen of the BY4742 collection.*** Includes ORF and gene names of all 329 hits from the screen. 4983 strains from the haploid knockout collection were assayed for growth in the presence of 300ppm MCHM on solid YPD media. While approximately 900 strains showed some noticeable growth differences on at least one replicate, the 329 strains showed clear growth differences on all 3 biological replicates, accounting for a stringent threshold for inclusion as a “hit” for the knocked out gene being required for MCHM resistance.

**Table S2: *Full results of significantly upregulated genes from RNAseq data in MCHM/YPD using BY4741 wildtype strain.*** Upregulated transcripts from the BY4741 RNAseq are listed in full. Included are the ORF names, as well as log2fold change and adjusted p-values supplied by the differential analysis. All genes are at least greater than a two-fold increase in expression in MCHM than YPD. Gene names, locations, and putative functions are concatenated to the rightmost columns of the table for further information on included ORFs.

**Table S3: *Full results of significantly downregulated genes from RNAseq data in MCHM/YPD using BY4741 wildtype strain.*** Downregulated transcripts from the BY4741 RNAseq are listed in full. Included are the ORF names, as well as log2fold change and adjusted p-values supplied by the differential analysis. All genes are at least greater than 2fold decreased in expression in MCHM than YPD. Gene names, locations, and putative functions are concatenated to the rightmost columns of the table for further information on included ORFs.

**Supplemental File 1: *Full GO terms for the genetic screen of BY4742 knockout collection.*** Excel file containing 3 tabs. Tab 1 gives DAVID GO term results for the biological process. Tab 2 gives DAVID GO term results for molecular function. Tab 3 gives DAVID GO term results for the cellular component.

**Supplemental File 2: *Full GO terms for significantly upregulated genes of BY4741 yeast treated with MCHM.*** Excel file containing 3 tabs. Tab 1 gives DAVID GO term results for the biological process. Tab 2 gives DAVID GO term results for molecular function. Tab 3 gives DAVID GO term results for the cellular component.

**Supplemental File 1: *Full GO terms for significantly downregulated genes of BY4741 yeast treated with MCHM.*** Excel file containing 3 tabs. Tab 1 gives DAVID GO term results for the biological process. Tab 2 gives DAVID GO term results for molecular function. Tab 3 gives DAVID GO term results for cellular component.

